# microRNA-seq of cartilage reveals an over-abundance of miR-140-3p which contains functional isomiRs

**DOI:** 10.1101/2020.01.29.925206

**Authors:** Steven Woods, Sarah Charlton, Kat Cheung, Yao Hao, Jamie Soul, Louise N Reynard, Natalie Crowe, Tracey E. Swingler, Andrew J. Skelton, Katarzyna A. Piróg, Colin G. Miles, Dimitra Tsompani, Robert M. Jackson, Tamas Dalmay, Ian M. Clark, Matt J. Barter, David A. Young

## Abstract

MiR-140 is selectively expressed in cartilage. Deletion of the entire miR-140 locus in mice results in growth retardation and early-onset osteoarthritis-like pathology, however the relative contribution of miR-140-5p or miR-140-3p to the phenotype remains to be determined. An unbiased small RNA sequencing approach identified miR-140-3p as significantly more abundant (>10-fold) than miR-140-5p in human cartilage. Analysis of these data identified multiple miR-140-3p isomiRs differing from the miRBase annotation at both the 5’ and 3’ end, with >99% having one of two seed sequences (5 ‘ bases 2-8). Canonical (miR-140-3p.2) and shifted (miR-140-3p.1) seed isomiRs were overexpressed in chondrocytes and transcriptomics performed to identify targets. miR-140-3p.1 and miR-140-3p.2 significantly down-regulated 694 and 238 genes respectively, of which only 162 genes were commonly down-regulated. IsomiR targets were validated using 3 ‘UTR luciferase assays. miR-140-3p.1 targets were enriched within up-regulated genes in rib chondrocytes of *Mir140*-null mice and within down-regulated genes during human chondrogenesis. Finally, through imputing the expression of miR-140 from the expression of the host gene *WWP2* in 124 previously published datasets, an inverse correlation with miR-140-3p.1 predicted targets was identified. Together these data suggest the novel seed containing isomiR miR-140-3p.1 is more functional than original consensus miR-140-3p seed containing isomiR.

## INTRODUCTION

MicroRNAs (miRNAs) are small non-coding RNAs that regulate gene expression. Mature miRNAs are processed from primary transcripts containing stem loops by cleavage mediated by Drosha and then Dicer proteins (1). The 5’ and 3’ sides of the stem loop can both give rise to mature miRNAs termed ‘-5p’ and ‘-3p’ respectively (2). These mature miRNAs are then loaded into the RNA-induced silencing complex (RISC) to mediate either mRNA degradation or translation inhibition of target mRNA through binding of the miRNA seed sequence (5’ bases 2-8) to the target 3’UTR (3). The level of complementarity between the seed and target mRNA is important for target repression. Binding to positions 2-7 of the miRNA may indicate target repression, however either an adenine binding to position 1 of the miRNA (7a1 targets), a match at position 8 (7m8 targets) or both (8mer targets) can improve target repression (4).

In recent years, small RNA (sRNA) sequencing (sRNA-seq) has identified additional miRNAs that do not perfectly align to the annotated mature miRNA, known as isomiRs (5). IsomiRs may be shorter or longer at the 5’ and/or 3’ end of the mature miRNA, and are not always templated to the genomic DNA. Variants at the 3’ end are quite common, although not thought to alter the miRNA target repertoire (6). Variations at the 5’ end are less common, but are of greater importance due to altered seed sequence, target repertoire and potentially the function of miRNAs.

The most studied miRNA in cartilage is miR-140, which produces miR-140-5p and miR-140-3p (7,8). Deletion of *Mir140* in mice leads to a skeletal phenotype including an OA-like disease. However, the contribution of the lack of miR-140-5p or miR-140-3p to the phenotype remains largely undetermined. Barter *et al*. pay particular attention to the role of miR-140-5p and suggests there are many genes under its control during chondrogenesis (9). miR-140-3p is less well studied and only a small number of -3p targets have been identified, which so far appear to have little relevance to cartilage and OA. Intriguingly, using small RNA sequencing (sRNA-seq) of human cartilage RNA we identified that miR-140-3p was high in abundance (>10-fold) compared to miR-140-5p (10).

Although there have been a number of studies to elucidate the role of miRNAs in cartilage and chondrogenesis, the role of isomiRs has been largely ignored. Here we show miR-140-3p isomiRs are abundantly expressed in cartilage and that these isomiRs are present in RISC. Using over expression followed by transcriptomic analysis we show two miR-140-3p isomiRs (miR-140-3p.1 and miR-140-3p.2) have largely discrete target repertoires. We validated a number of common and discrete targets for each isomiR using a luciferase reporter system. We present evidence that miR-140-3p.1, which is not currently annotated in miRBase, may play roles in human chondrogenesis, mouse cartilage and in multiple other skeletal tissues.

## MATERIAL AND METHODS

### Analysis of cartilage sRNA-seq

sRNA-seq was analysed as previously described (10). 5’ isomiRs were defined using the following criteria: 1) loss or gain of 1 or more nucleotides, using the mature miRBase (11) sequences as a reference; 2) a read count >100, and 3) a read count > 5% of the mature miRBase reference sequence for that miRNA.

### Human articular chondrocyte isolation, culture and transfection

Human articular chondrocyte isolation from knee cartilage was performed as previously described (9). Tissue was donated by patients with diagnosed osteoarthritis and undergoing joint replacement surgery, with informed consent and ethics committee approval. Briefly, macroscopically normal cartilage was removed from the subchondral bone and dissected into ~1mm pieces using scalpel and forceps. Enzymatic digestion was performed using trypsin and then collagenase overnight at 37°C. Chondrocytes were then grown to confluence and plated into 6 well plates for miRNA/isomiR mimic transfection. 100nM miRNA mimic and control were transfected into HAC using DharmaFECT 1 transfection reagent (Dharmacon, Horizon Discovery, Cambridge, UK) according to the manufacturer’s protocol and essentially as previously described (9). 48h post-transfection HAC were lysed and RNA harvested using Qiagen miRNeasy kit (Qiagen, Crawley, UK). Custom miRNA mimics with each isomiR sequence were purchased from Dharmacon.

### Microarray and identification of targets

Microarray was performed by Cardiff University, Central Biotechnology Services, using Illumina whole-genome expression array Human HT-12 V4 (Illumina, Saffron Walden, U.K.). Normalization of the quantified signals (background corrected) was performed using the global Lowess regression algorithm. Expression analysis was performed in R/bioconductor using the limma package (12). PCA and heat map were generated using ClustVis (13). PCA analysis was performed on all unique genes detected by the microarray analysis. Heat map is based on the top 650 significant genes for each comparison (miC vs. mR-140-3p.1 or miC vs. miR-140-3p.2), which after removal of duplicates gave 882 genes. TargetScanHuman (version 7.2 (14)) was used to predict targets of miR-140-3p.1 and miR-140-3p.2. In order to demonstrate discrete functions we investigated unique targets for each isomiR (3p.2 and 5p targets were removed from 3p.1 analysis; 3p.1 and 5p targets were removed from 3p.1 analysis; 3p.1 and 3p.2 targets were removed from 5p analysis). Genes were selected for luciferase validation based on both altered expression in miR-140-3p.1 or miR-140-3p.2 transfected cells versus control and on expression in miR-140-3p.1 transfected cells versus expression in miR-140-3p.2 transfected cells.

### Target enrichment and pathway analysis

Sylamer analysis was performed on ordered gene lists from most downregulated to most upregulated (including genes whose expression did not significantly change) (15). Target enrichment (%targets) was calculated by dividing the total number of predicted targets (TargetScan 7.2) that significantly increased or decreased by the total number of genes that significantly increased or decreased, multiplied by 100. Cumulative fraction plots were generated using ordered gene list from down-regulated to most up-regulated (regardless of significance) and predicted targets from TargetScan 7.2. Pathway analysis was performed using Database for Annotation, Visualization and Integrated Discovery (DAVID) v6.7(16).

### Q-RT-PCR for miR-140-3p.1 and miR-140-3p.2

Custom miR-140-3p.1 and miR-140-3p.2 assays (Exiqon) were used to detect expression of miR-140-3p.1 and miR-140-3p.2. Assays were validated using spike-in of miR-140-3p.1 and miR-140-3p.2 mimic (Dharmacon).

### 3’UTR luciferase reporter construction and assay

In-Fusion® cloning (Clontech, Takara Bio Europe SAS, Saint-Germain-en-Laye, France) of selected 3’UTRs into the pmirGLO vector (Promega, Southampton, UK) was used to generate 3’UTR luciferase reporters essentially as previously described (9). SW1353 chondrosarcoma cells were seeded and cultured to reach ~50% confluence after 24 h (9). miRNA (100 nM) were transfected using DharmaFECT 1 transfection reagent, reporter plasmids (500 ng/ml) were transfected using FugeneHD (Promega). 24h after transfection luciferase levels determined using Promega dual luciferase assay and GloMax plate reader (Promega).

### Generation of miR-140^−/−^ mice

All animal experiments were performed under licenses granted from the Home Office (United Kingdom) in accordance with the guidelines and regulations for the care and use of laboratory animals outlined by the Animals (Scientific Procedures) Act 1986 according to Directive 2010/63/EU of the European Parliament, and conducted according to protocols approved by the Animal Ethics Committee of Newcastle University and the Home Office, United Kingdom. Post-generation breeding and subsequent phenotyping were performed under licences PPL60/4525 and P8A8B649A. CRISPR/Cas9 guide RNAs (crRNA) were designed using CHOPCHOP. crRNA linked with TRACR (sgRNA) were amplified by PCR with a pLKO vector (Addgene_52628) as template, using the primer T7 TRACR R (Supplementary Table 10) and a 5’ PCR primer that included a T7 sequence and the crRNA. This was converted to RNA using the MEGAshortscript T7 kit (Fisher Scientific). sgRNA (50ng/ml each) were mixed with recombinant Cas9 (ToolGen, CamBioScience Limited, Cambridge, UK) and injected into the cytoplasm of donor mouse zygotes and transferred into recipient foster mothers, all essentially as previously described (17–19). The mixed C57BL/6 and CBA/ca F_0_ mice were backcrossed onto C57BL/6J and heterozygous animals crossed three times to eventually generate wild-type and null lines used in purification of rib chondrocytes.

Genotype was confirmed by ear-notch PCR and Sanger sequencing (Supplementary Figure 3).

### Rib chondrocyte isolation and RNA-seq

Primary mouse costal chondrocytes were isolated from 7-day-old miR-140^−/−^ and WT mice using collagenase digestion, essentially as previously described (20). Total RNA, including miRNA was isolated using the miRVana miRNA Isolation Kit (with phenol)(Fisher Scientific). Sequencing libraries were prepared from 500 ng of purified total RNA using the Illumina TruSeq Stranded mRNA sample preparation kit according to the manufacturer’s protocol, and sequenced on Illumina NextSeq500. Each sample provided >12 million single-end 75-bp sequencing reads. Sequenced reads were mapped to the mm10 transcriptome using Salmon (21). Batch effects were estimated using RUVseq (22) and incorporated into the differential expression analysis performed using DESeq2 (23). PCA and heat map were generated using ClustVis (13).

### SkeletalVis analysis

Gene expression responses within SkeletalVis were filtered for human and mouse data sets and where *WWP2* expression significantly changed (adjusted p-value is <0.05), leaving 124 experimental comparisons. The change in WWP2 expression was plotted against the average fold change of predicted (TargetScan7.2) miRNA targets for each experimental comparison. Correlation (R^2^) and regression analysis for all 124 experimental comparisons was calculated using the data analysis add-in in Microsoft Excel. Target enrichment (%targets) was calculated by dividing the total number of predicted targets (TargetScan 7.2) that significantly increased or decreased by the total number of genes that significantly increased or decreased, multiplied by 100, for each of the 124 experimental comparisons. miR-140-5p, miR-140-3p.1 and miR-140-3p.2 target enrichment within up, no change or down-regulated genes was then plotted against log2 FC for *WWP2*. Cumulative mean enrichment (from studies where *WWP2* is most up- or down-regulated compared to studies where *WWP2* changes least, were plotted to demonstrate trend. Correlation between miR-140-5p, miR-140-3p.1 and miR-140-3p.2 target enrichment was plotted by first calculating the relative enrichment in up vs down-regulated genes for each experimental comparison (enrichment in up/enrichment in down). This relative enrichment for each of miR-140-5p, miR-140-3p.1 and miR-140-3p.2 was plotted against each other. Experimental comparisons where *WWP2* decreased or increased were shown in different colours as indicated in figure legend. A list of randomly generated genes was analysed alongside miR-140-5p, miR-140-3p.1 and miR-140-3p.2.

## RESULTS

### RNA-Seq of articular cartilage identifies an overabundance of miR-140-3p and miR-140-3p isomiRs

We performed sRNA-seq of human cartilage RNA and identified novel cartilage specific miRNAs (10). Our cartilage sRNA-seq aligned to 990 miRNAs annotated in miRbase (11), 704 of which contained at least one additional type of potential isomiR (Table S1). As 5’ isomiRs are most likely to alter function, we focused further sRNA-seq analysis on miRNAs with 5’isomiRs. Using thresholds of >100 reads and >5% of total reads for that miRNA, we identified 5’isomiR sequences for 29 miRNAs, 26 of which have a single addition or deletion at the 5’ nucleotide, while miR-3074-5p, miR-455-3p and let-7b-3p all have two additional 5’ nucleotides (Supplementary figure 1). Although the isomiR for miR-1246 is just under our read threshold (read count of 99), it appears to have four additional 5’ nucleotides compared to the miRBase annotation (Supplementary figure 1).

MiR-140-3p and its isomiRs account for more than half of all sequencing reads in our cartilage sRNA-seq (Figure 1A). IsomiRs can be sub-divided into several categories, essentially either templated or non-templated with either 5’ or 3’ modifications (24). miR-140-3p isomiRs are predominantly 3’ additions and mixed type isomiRs, with only 5% aligning to the original miRBase annotation (Figure 1B). More than 99% of all miR-140-3p isomiRs result in one of two seed sequences; ACCACAG as annotated in miRBase, and CCACAGG, which is shifted by −1 nucleotide (Figure 1C). Twenty-five percent of reads for miR-140-3p were perfectly templated to DNA (Figure 1D), the remainder contained one or more non-templated nucleotides, predominantly non-templated 3’ adenine additions (Figure 1E), which were observed in sequencing reads containing both of the seed sequences (Figure 1F and G). The most detected isomiR with each seed are termed miR-140-3p.1 (ACCACAGGGTAGAACCACGGAC, seed CCACAGG) and miR-140-3p.2 (TACCACAGGGTAGAACCACGGA seed: ACCACAG), respectively (Figure 1H). Expression of miR-140-3p.1 and miR-140-3p.2 in cartilage was validated using qRT-PCR with isomiR selective assays (Figure 1I).

**Figure 1.**
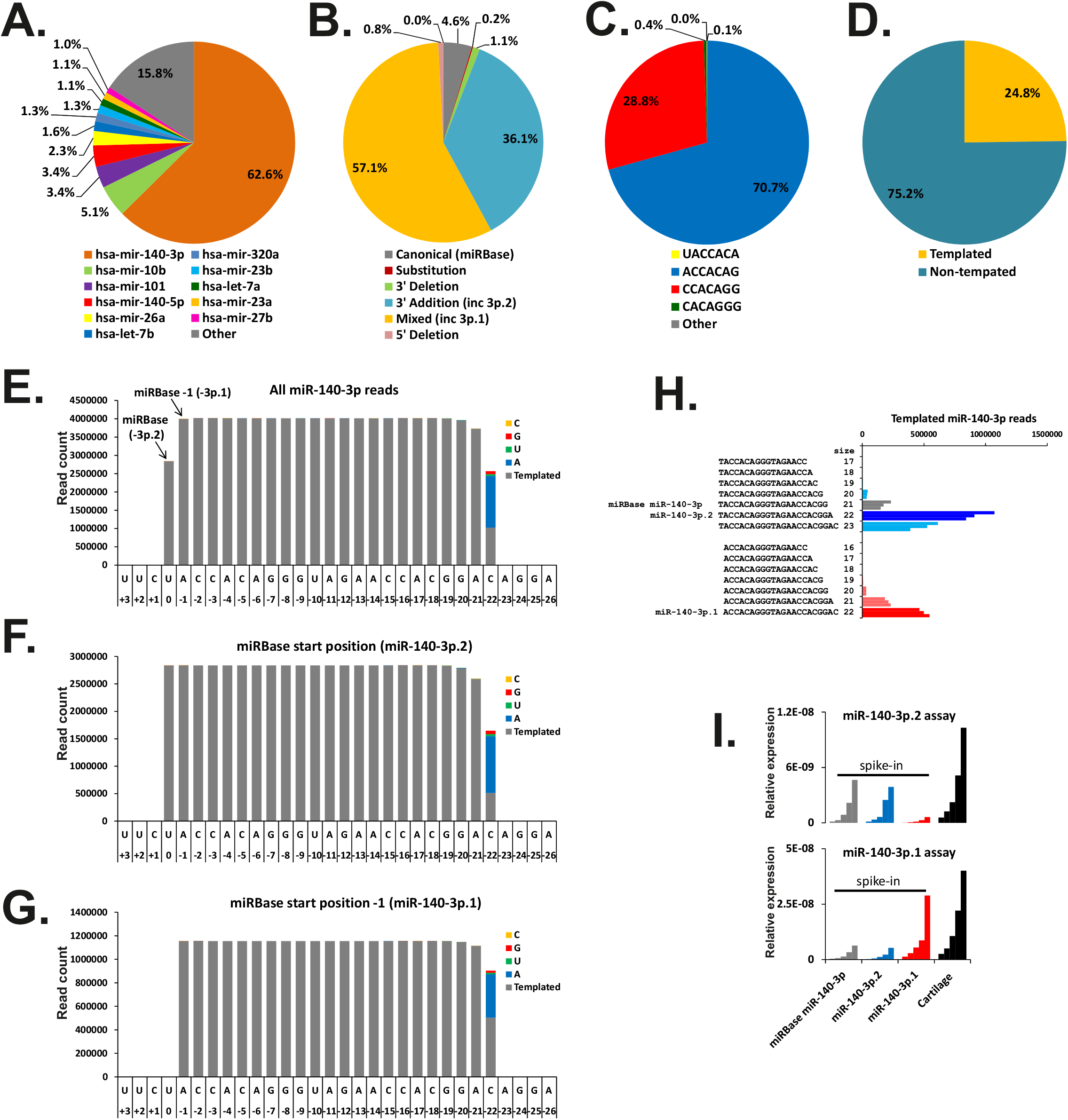
miR-140-3p isomiRs are abundantly expressed in cartilage. (A) Pie chart illustrating the relative abundance of all miRNAs expressed in cartilage, data combines isomiR and canonical reads for each miRNA. (B) Pie chart illustrating the isomiR type for all reads aligned to miR-140-3p. (C) Pie chart illustrating seed sequences for all reads aligned to miR-140-3p, >99% of reads have one of two seed sequences; ACCACAG (blue) and CCACAGG (red). (D) Pie chart of the percentage of templated and non-templated reads for miR-140-3p within cartilage. (E) Histogram of all sequencing reads aligned to miR-140-3p in cartilage. (F) Histogram of sequencing reads aligned to miR-140-3p, whose 5’ end is as indicated in miRBase. (G) Histogram of sequencing reads aligned to miR-140-3p, whose 5’ end is one nucleotide shorter than indicated in miRBase. (H) Bar chart indicting the sequences and number of reads that contribute to each of the two seed sequences. Individual bars represent total read number from 3 separate individuals. (I) qRT-PCR for miR-140-3p.1 and miR-140-3p.2. Assays designed to detect each isomiR are able to distinguish between miR-140-3p.1 and miR-140-3p.2 spike-ins (2 fold serial dilution) and detect high expression of each isomiR in cartilage (2 fold serial dilution).

Analysis of other published sRNA-seq data identified the presence of miR-140-3p.1 and miR-140-3p.2 in other human tissue types, although expressed at a lower level than in cartilage (Supplementary figure 2) (25–27). As miRNA bound to Argonaute (AGO) indicates the ability to repress targets (28), we analysed sRNA-seq data following immunoprecipitation of AGO proteins (29). Indeed both miR-140-3p.1 and miR-140-3p.2 were present, indicating each can be loaded into RNA-induced silencing complex (RISC; Supplementary figure 2), and have the potential to target transcripts.

### miR-140-3p.1 and miR-140-3p.2 are predicted to have distinct targets

As miRNAs generally target mRNAs through interaction of their seed sequence (nucleotides 2-7), the seed sequences for miR-140-3p.1 and miR-140-3p.2 differ (CCACAGG and ACCACAG, respectively; Figure 2A). Thus each miR-140-3p isomiR has different preferences for seed binding sites (Figure 2A). Indeed, target prediction analysis for miR-140-3p.1 and miR-140-3p.2 by TargetScan 7.2 (14) (conserved) identifies only 133 shared targets, a small number compared to the total number of predicted targets (665 and 496 for miR-140-3p.1 and miR-140-3p.2, respectively, Figure 2B, Table S2). The majority of these targets also differ from predicted miR-140-5p targets (Figure 2B, Table S2).

**Figure 2.**
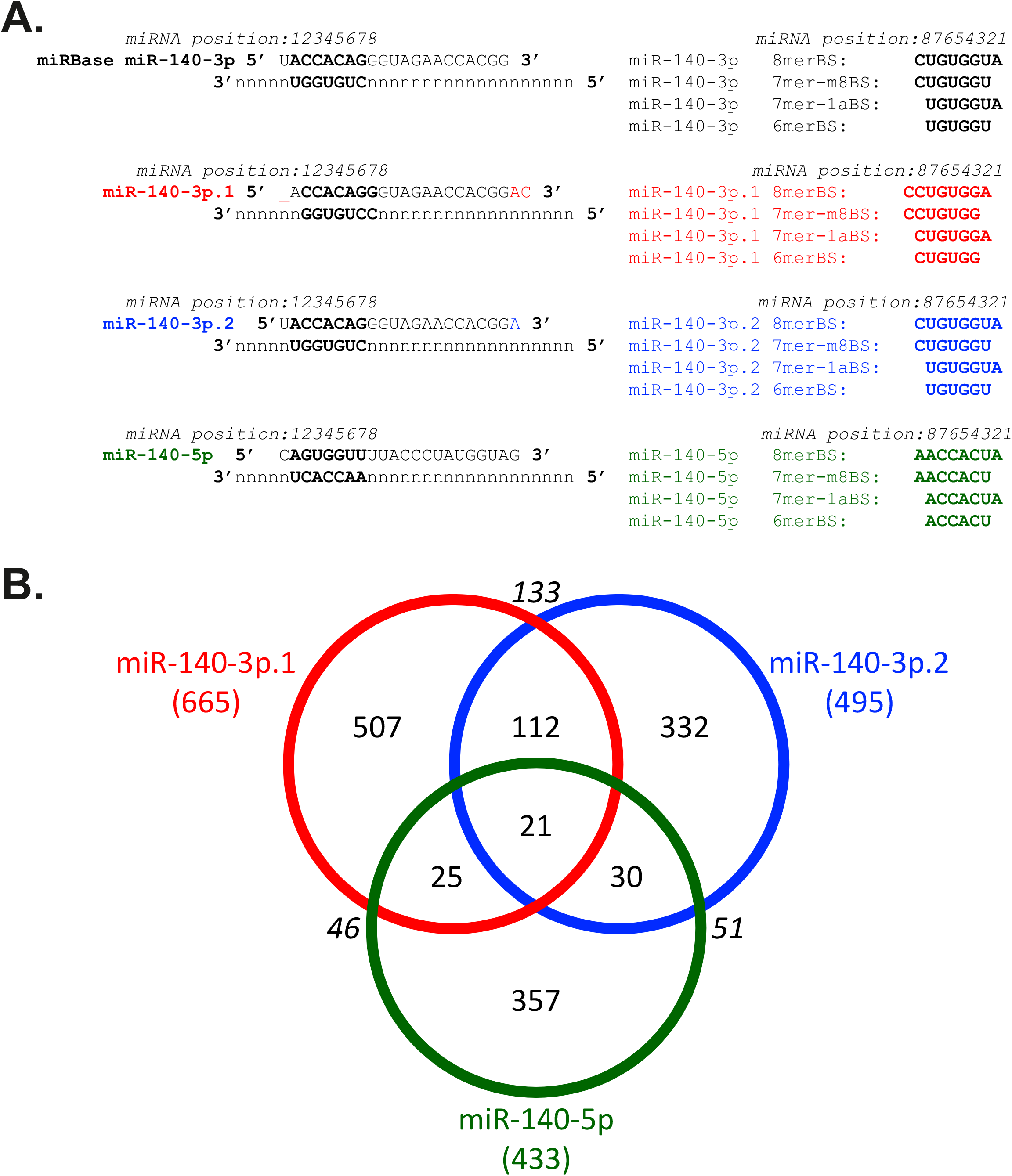
miR-140-3p.1 and miR-140-3p.2 are predicted to have distinct targets. (A) Schematic illustrating the sequences of miRBase miR-140-3p, miR-140-3p.1, miR-140-3p.2 and miR-140-5p and their potential binding sites within target mRNAs using the TargetScan categorisation. (B) Venn diagram illustrating little cross over of targets predicted by TargetScan 7.2 for miR-140-3p.1, miR-140-3p.2 and miR-140-5p.

### Identification and validation of miR-140-3p.1 and miR-140-3p.2 targets in human chondrocytes

Next, we used overexpression of miR-140-3p.1 and miR-140-3p.2 in primary human chondrocytes followed by genome-wide transcriptomics to experimentally identify similarities and differences in their target repertoire. Overexpression of miR-140-3p.1 and miR-140-3p.2 was validated using isomiR-specific qRT-PCR (Figure 3A). Principal component analysis (PCA) and heat map illustrated clustering of biological replicates and differences between miR-140-3p.1 and miR-140-3p.2 in transfected chondrocytes (Figure 3B and C). 693 and 237 genes were significantly (adj.P.Val<0.05) down-regulated by miR-140-3p.1 and miR-140-3p.2 respectively, however only 161 genes were commonly down-regulated by both isomiRs (compared to control;Figure 4D, Table S3, Table S4). Pathway analysis of repressed genes identified enrichment terms associated with ‘phosphoprotein’ for miR-140-3p.1 (Table S5). Direct comparison of miR-140-3p.1 transfected with miR-140-3p.2 transfected cells revealed 237 genes were significantly different between the two isomiRs (Table S3, Table S4). Pathway analysis of the 237 genes differentially expressed between the two isomiRs identified an enrichment of triple helix collagen genes (*COL2A1, COL4A1, COL6A3, COL11A1, COL11A2*; p=0.0004, adjp=0.066; Table S5), indicating the sequence difference between the two isomiRs is functionally important for chondrocytes/cartilage.

**Figure 3.**
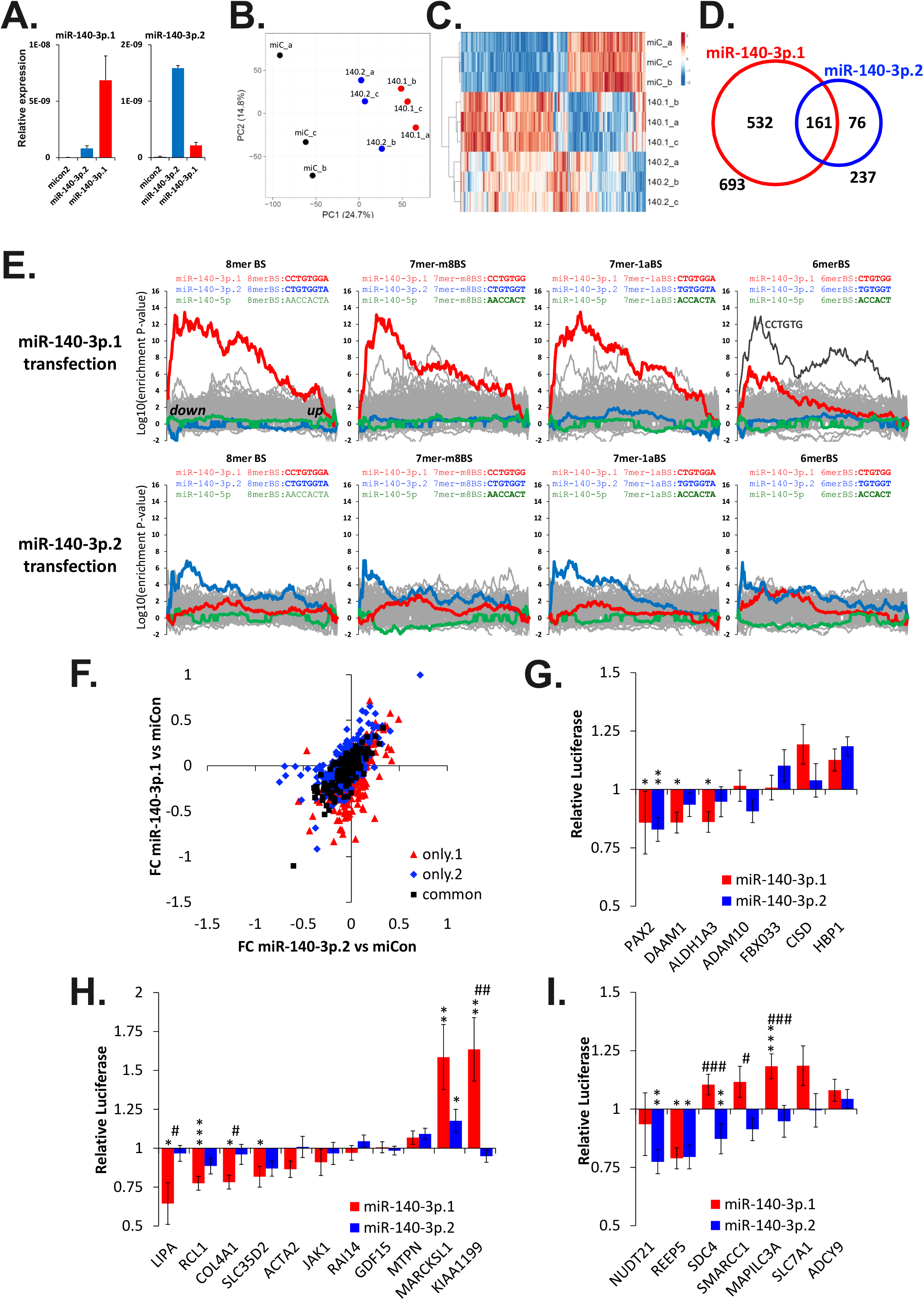
Identification and validation of miR-140-3p.1 and miR-140-3p.2 targets in human chondrocytes. (A) qRT-PCR for miR-140-3p.1 and miR-140-3p.2 following transfection of either control mimic, miR-140-3p.1 mimic or miR-140-3p.2 mimic for 48h. (B) PCA of whole genome micro-array following transfection of mimics described in (A), (140.1= miR-140-3p.1 and 140.2=miR-140-3p.2). (C) Heat map of whole genome micro-array following transfection of mimics as described in (A) (140.1= miR-140-3p.1 and 140.2=miR-140-3p.2). (D) Venn diagram indicating the number of genes down-regulated following overexpression of miR-140-3p.1 and miR-140-3p.2. (E) Enrichment analysis using Sylamer for 3’UTR mRNA motifs complementary to either miR-140-3p.1 or miR-140-3p.2 seed sequences in genes whose expression decreased following transfection of either miR-140-3p mimic, miR-140-3p.1 mimic or miR-140-3p.2 mimic. (F) Gene expression changes of miR-140-3p.1 (only.1;red), miR-140-3p.2 (only.2;blue) or miR-140-3p.1 and miR-140-3p.2 (common;green) predicted targets following transfection of either miR-140-3p.1 mimic or miR-140-3p.2 mimic. (G-I) 3’ UTR luciferase reporters for miR-140-3p.1 and miR-140-3p.2 putative targets. *, ** or *** indicate level of significance (p<0.05, 0.01 or 0.001, respectively) of either miR-140-3p.1 or miR-140-3p.2 to control. # indicates level of significance (p<0.05, 0.01 or 0.001, respectively) between miR-140-3p.1 and miR-140-3p.2.

**Figure 4.**
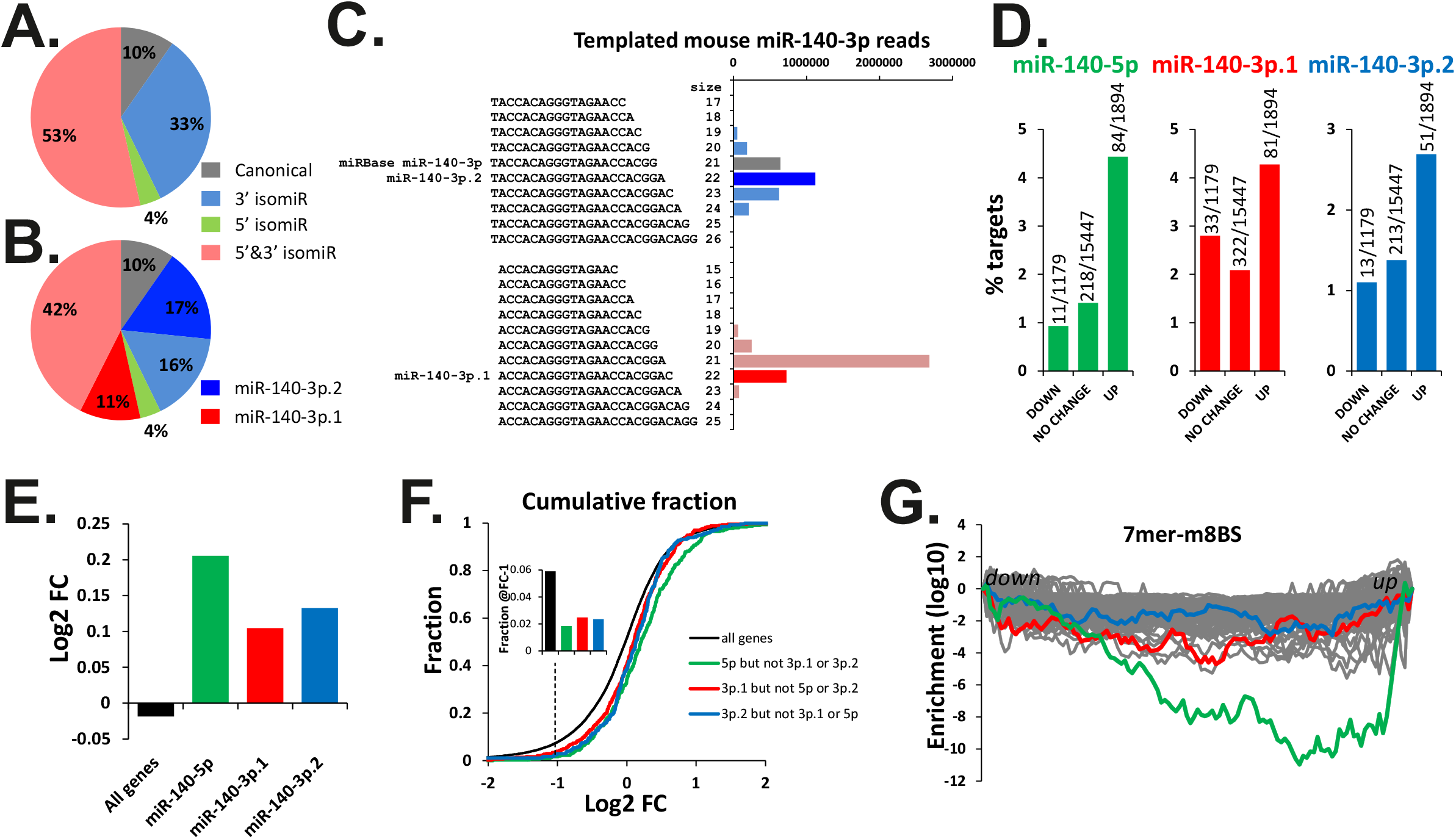
Evidence for functional miR-140-3p.1 and miR-140-3p.2 in mouse cartilage. (A) Pie chart illustrating the fraction of miR-140-3p reads with canonical 5’ and 3’ ends (grey), canonical 5’ and either shorter or longer 3’ (blue), canonical 3’ and either shorter or longer 5’ (green), either shorter or longer 5’ and either shorter or longer 3’ (red). (B) As in (A) with miR-140-3p.1 and miR-140-3p.2 indicated in dark red and dark blue respectively. (C) Bar chart indicting number of reads that contribute to (B). (D) % of genes that are predicted targets of either a miR-140-5p (green), miR-140-3p.1 (red) or miR-140-3p.2 (blue), in genes whose expression decrease, does not change or increase following KO of miR-140 in mice. (E) Median fold change (FC) of all genes that are predicted targets of either a miR-140-5p (green), miR-140-3p.1 (red) or miR-140-3p.2 (blue) following KO of miR-140 in mice. (F) Cumulative fraction plot of all genes (black), miR-140-5p (green), miR-140-3p.1 (red) or miR-140-3p.2 (blue) predicted targets, for genes ordered from most decreased expression to most increased expression in Mir140 null mice. Inset: Fraction at a cut-off of log2FC −1. (G) Sylamer analysis for Mir140 null mice; the 100 most enriched 7mers are shown with 7m8 binding sites for miR-140-5p (green), miR-140-3p.1 (red) and miR-140-3p.2 (blue) highlighted.

Sylamer, a non-bias motif analysis tool, was used to investigate miRNA binding site enrichment within downregulated 3’UTRs (15). Transfection of either miR-140-3p.1 or miR-140-3p.2, showed specific enrichment for 8mer, 7m8, 7a1 and 6mer seed binding sites of the transfected isomiR. Interestingly, a greater enrichment was observed for miR-140-3p.1 (non-canonical seed) binding sites than miR-140-3p.2 (canonical seed) binding sites (Figure 3E). Predicted targets (identified using TargetScan 7.2) that are unique to either miR-140-3p.1 or miR-140-3p.2 were downregulated following overexpression of either miR-140-3p.1 or miR-140-3p.2 respectively, but not downregulated by the reciprocal isomiR (Figure 3F), suggesting both isomiRs can repress specific transcripts in human chondrocytes.

Unique and common targets were then selected for 3’UTR luciferase validation (Figure 3G-I). As expected *LIPA*, *RCL1*, *COL4A1* and *SLC35D2* were validated as miR-140-3p.1 targets; *NUDT21*, *REEP5* and *SDC4* were validated as miR-140-3p.2 targets and *PAX2* validated as a target of both miR-140-3p.1 and miR-140-3p.2. Interestingly, the *REEP5* 3’UTR luciferase constructs was also repressed by miR-140-3p.1, although not a predicted target of miR-140-3p.1. Conversely, *DAAM1* and *ALDH1A3* are predicted targets of both miR-140-3p.1 and miR-140-3p.2, however significant repression was only observed for miR-140-3p.1 and not for miR-140-3p.2. Contrary to the repression role of miRNAs in gene regulation, miR-140-3p.1 increased luciferase for the *MARCKSL1*, *KIAA1199* and *MAPILC3A* 3’UTR reporters.

### miR-140-3p isomiRs are functional in mouse cartilage

Analysis of mouse sRNA-Seq data revealed that both miR-140-3p.1 and miR-140-3p.2 are present and more abundant than miR-140-5p (30) (Figure 4A-C), indicating miR-140-3p isomiRs are conserved across species. We and others have generated *Mir140* null mice, which lack all miRNAs and isomiRs produced from the *Mir140* locus (Supplementary figure 3). Whole genome RNAseq data of rib cartilage from these mice identified 1894 and 1179 genes that were significantly up and down-regulated respectively (Table S6, Table S7). Pathway analysis of upregulated genes following knock out (KO) of *Mir140* identified terms associated with ‘phosphoprotein’ (Table S8), which is consistent with pathway analysis of genes decreased following miR-140-3p.1 overexpression in human chondrocytes (Table S5). miR-140-5p, miR-140-3p.1 and miR-140-3p.2 predicted targets (identified using TargetScan 7.2) were enriched within upregulated genes (Figure 4D), and their average expression increased following KO of the *Mir140* locus (Figure 4E), indicating a loss of target repression. Furthermore the cumulative fraction plot indicates the fraction of targets that decrease following KO of the *Mir140* locus is lower than expected (Figure 4F). Non-biased Sylamer analysis also identified an enrichment of miR-140-5p, miR-140-3p.1 and miR-140-3p.2 7m8 seed binding sites within upregulated genes; miR-140-5p displayed the greatest enrichment, followed by miR-140-3p.1 and then miR-140-3p.2 (Figure 4G).

More than 99% of miRNAs produced from the murine *Mir140* locus share seed binding sites with either miR-140-5p, miR-140-3p.1 or miR-140-3p.2; of these miR-140-5p has the lowest read count, yet has the largest contribution to target repression (Figure 4). We used Sylamer to investigate the possibility that other lowly expressed sequences produced from the *Mir140* stem loop may contribute to target repression (Supplementary figure 4A). This analysis indicates miR-140-5p followed by miR-140-3p.1 have the largest contribution to target repression in mir140 KO mice (Supplementary figure 4B), and that there are no lowly expressed *Mir140*-derived miRNAs that have a major contribution to target repression in mice (Supplementary figure 4B).

### miR-140-3p.1 targets are enriched within down-regulated genes during MSC chondrogenesis

miR-140 is highly up-regulated during mesenchymal stem cell (MSC) chondrogenesis, with miR-140-5p predicted targets highly enriched within the down-regulated genes (9). Here we show through re-analysis of this transcriptomic dataset (Figure 5A) that expression of predicted targets of both miR-140-3p.1 and miR-140-3p.2 are decreased during chondrogenesis. Similar to miR-140-5p predicted targets, the average expression of miR-140-3p.1 and miR-140-3p.2 predicted targets decreased during chondrogenesis, with unique miR-140-3p.1 targets being more repressed than unique miR-140-3p.2 predicted targets (p=0.07) (Figure 5B). Sylamer analysis indicated enrichment for miR-140-5p and miR-140-3p.1, but minimal enrichment for miR-140-3p.2 (Figure 5C). Consistent with *Mir140* null mouse data, the enrichment of miR-140-5p was greater than for both miR-140-3p.1 and miR-140-3p.2 in genes whose expression decreased during human MSC chondrogenesis (Figure 5C).

**Figure 5.**
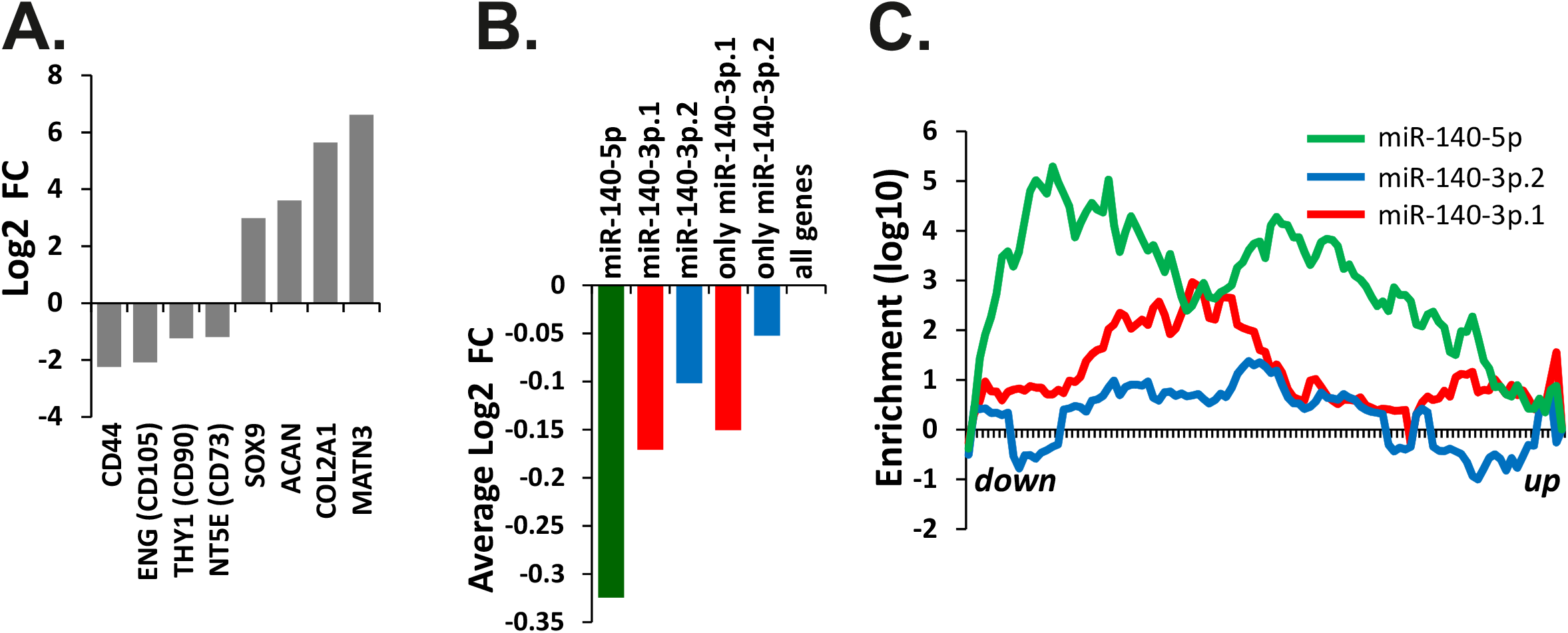
Evidence for miR-140-3p.1 and miR-140-3p.2 function during human MSC chondrogenesis. Re-analysis of data from Barter *et al* [10] for human MSC differentiation to cartilage. (A) Changes in gene expression for eight MSC and chondrogenic markers. (B) Mean fold change of all genes that are predicted targets of either a miR-140-5p (green), miR-140-3p.1 (red) or miR-140-3p.2 (blue) during MSC chondrogenesis. (C) Sylamer analysis for a miR-140-5p (green), miR-140-3p.1 (red) and miR-140-3p.2 (blue) 7m8 seed sequences.

### miR-140-5p and miR-140-3p.1 predicted targets inversely correlate with *WWP2* in multiple skeletal tissues

Having established miR-140-5p, miR-140-3p.1 and miR-140-3p.2 targets are regulated during human MSC chondrogenesis and in mouse rib chondrocyte development, we next looked for evidence of an inverse correlation between miR-140 and target gene expression in other skeletal tissues. Whole genome transcriptomic datasets of skeletal tissues following various perturbations have been collated within SkeletalVis, a data portal for cross-species skeletal transcriptomics data (31). Although the majority of these studies do not directly assess miRNA expression, miRNAs that are located in the introns of protein coding genes, including miR-140, and are frequently co-regulated with their host gene (32). A number of studies have utilised host-gene expression to predict the gene targets of intronic miRNAs through a consistent negative correlation in expression (33,34). miR-140 is located in intron 16 of *WWP2*, with expression of the miR and an abundant isoform of *WWP2* (transcript variant 2) controlled by a common promoter (35). *WWP2* was therefore used as a surrogate for miR-140 expression.

Of the 779 skeletal gene expression responses within SkeletalVis, 124 contained a significant alteration (adjusted p-value <0.05) in *WWP2* expression. The change in *WWP2* expression (as a surrogate for mir140 expression) was plotted against the average fold change of predicted miRNA targets for each study. Using regression analysis we identified a significant inverse relationship between *WWP2* and miR-140-5p (Figure 6A; r^2^=0.38, p=3.08×10^−14^) and miR-140-3p.1, predicted targets (Figure 6B; r^2^=0.16, p=5.46×10^−6^). There was also minimal but significant inverse relationship between *WWP2* and miR-140-3p.2 predicted targets (Figure 6C; r^2^=0.05, p=0.012). Thus, where *WWP2* expression changes, expression of predicted miR-140 targets have a tendency to change in the opposite direction. Next we determined if there was enrichment for miR-140 predicted targets in genes whose expression level changes reciprocally to *WWP2.* Indeed, within studies where *WWP2* expression increased there was generally a larger percentage of predicted miR-140-5p and miR-140-3p.1 targets within down-regulated genes than up-regulated genes (Figure 6E and F). However, when *WWP2* expression decreased, only miR-140-5p predicted targets were enriched in the up-regulated genes (Figure 6E). There was minimal enrichment of miR-140-3p.2 targets (Figure 6G).

**Figure 6.**
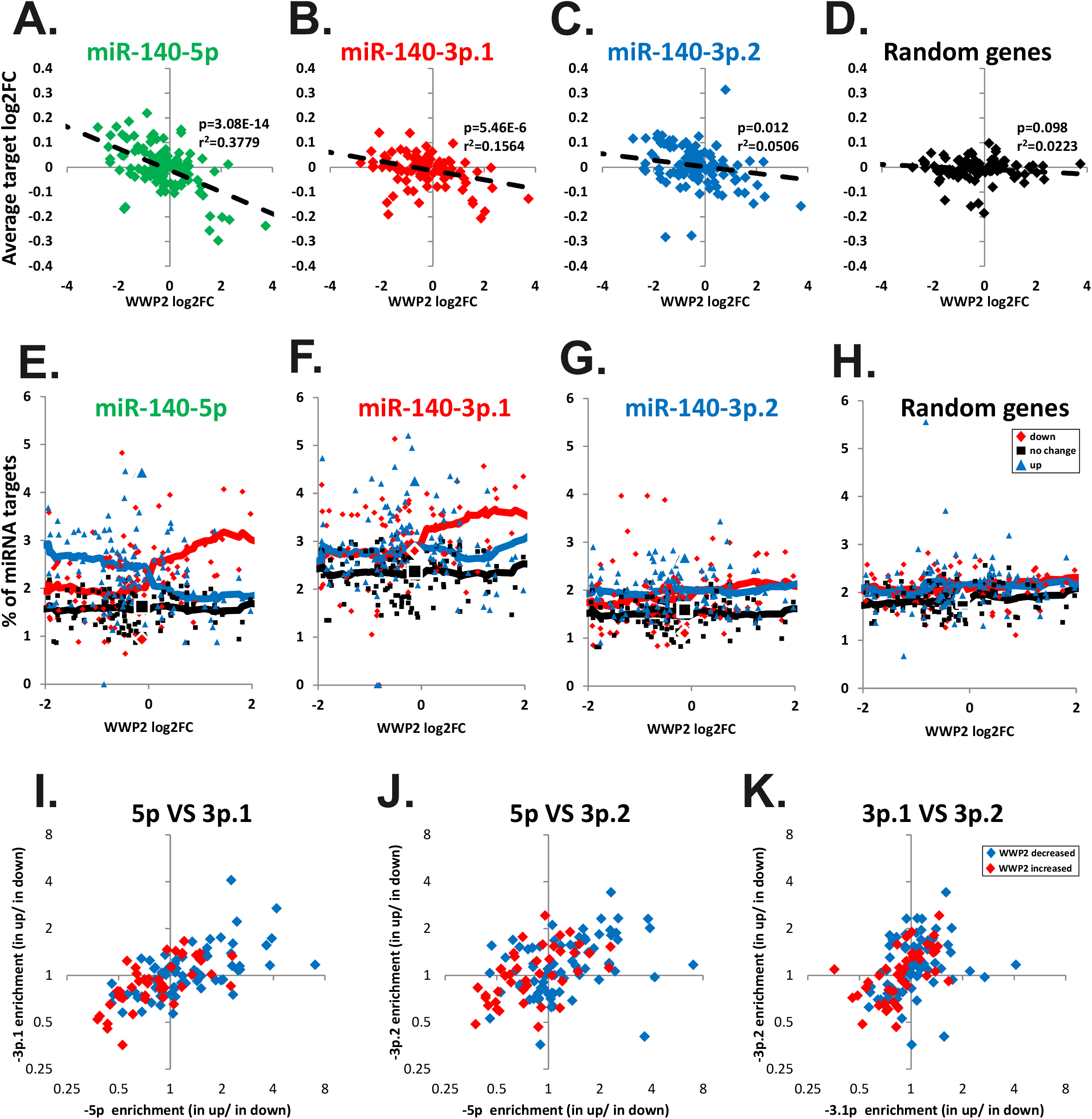
Correlation of mir140 host gene WWP2 with mir140 target genes in SkeletalVis transcriptomic datasets. (A-D) Correlation analysis of WWP2 and predicted miRNA target expression in 124 pre-existing transcriptome-wide comparisons (both human and mouse). Scatter graph of WWP2 log2FC plotted against the average log2FC for genes predicted to be either a target of miR-140-5p (A), miR-140-3p.1 (B), miR-140-3p.2 (C) or 500 random genes (D), p-value calculated from linear regression analysis. (E-H) Enrichment analysis of miR-140-5p (E), miR-140-3p.1 (F) or miR-140-3p.2 (G) predicted targets and 500 random genes (H) in data sets where WWP2 expression significantly changes. Data is plotted as percentage of targets within up-regulated, no change or down-regulated transcripts against log2FC WWP2. Large icons represent enrichment of miR-140-5p, miR-140-3p.1 and miR-140-3p.2 predicted targets in up- and down-regulated genes within rib cartilage of MIR140 null mice, where Wwp2 expression does not change. (I-K) Correlation of miR-140-5p and miR-140-3p.1 target enrichment. In studies where WWP2 significantly decreased (blue) and increased (red). The enrichment of miR-140-5p and miR-140-3p.1 and miR-140-3p.2 targets in significantly down and up-regulated genes was calculated. From this the relative enrichment in up vs down-regulated genes was calculated (enrichment in up/enrichment in down). The relative enrichment for each miRNAs targets were plotted against each other.

To determine if miR-140-5p and miR-140-3p isomiR targets are regulated in a similar way in each study we correlated relative enrichment (enrichment in upregulated divided by enrichment in down-regulated genes) for miR-140-5p, miR-140-3p.1 and miR-140-3p.2 predicted targets. Enrichment observed for miR-140-5p targets correlated with enrichment for miR-140-3p.1 targets (Figure 6I). Studies where miR-140-5p and miR-140-3p.1 predicted targets display a greater enrichment in up-regulated genes than in down-regulated genes (enrichment in up divided by enrichment down >1), were mostly studies where *WWP2* decreased (Figure 6I). Likewise in studies where miR-140-5p and miR-140-3p.1 predicted targets display a greater enrichment in down-regulated genes than up-regulated genes (ratio of up:down-regulated <1), were mostly studies where *WWP2* increased (Figure 6I). These data indicate that when *WWP2* expression increases, both miR-140-5p and miR-140-3p.1 predicted targets are repressed. Furthermore, they also indicate that miR-140-3p.2, essentially the canonical form of miR-140-3p, has a less significant biological effect on gene expression than miR-140-5p or miR-140-3p.1.

To further refine prediction of putative direct miR-140 targets, we investigated the overlap between four criteria; A) TargetScan predicted targets (Figure 2), B) genes down-regulated following overexpression in human chondrocytes (Figure 3), C) genes upregulated in mice lacking the *Mir140* locus (Figure 4) and D) genes up-regulated during MSC chondrogenesis (Figure 5). There were 5, 4 and 0 genes matching all of these criteria for miR-140-5p, miR-140-3p.1 and miR-140-3p.2 respectively (Supplementary Figure 5A, Table S9). *ABCA1* was common to both miR-140-5p and miR-140-3p.1. Furthermore, *ABCA1* was inversely correlated with *WWP2* across multiple skeletal data sets, with the inverse correlation being strongest in cartilage (Supplementary Figure 5B), where miR-140 expression is highest.

## DISCUSSION

IsomiRs of miR-140-3p have previously been detected in chondrocytes (36), breast cancer cell lines (37) and endometrial tissue (38), and are now recognised in TargetScan 7.2 (39). In TargetScan and this study, miR-140-3p.2 refers to the original miRBase annotation and miR-140-3p.1 refers to the newly identified isomiR. MiRBase does not currently include specific annotation for isomiRs. In addition to miR-140-3p isomiRs, we detected isomiRs for a number of other miRNAs in cartilage, including miR-455, which is also now recognised in TargetScan 7.2 (39). Sequence analysis of the miR-1246 isomiR, suggests it may in fact be miR-1290. According to miRBase these two miRNAs are transcribed from two separate loci on chromosome 2 and 1, respectively. However, Mazieres *et al.* (40), suggest miR-1246 and miR-1290 are processed from a common transcript (RNU2), which explains their similar sequence and expression pattern during chondrogenesis (9). Many of the other detected cartilage isomiRs can be transcribed from more than one genomic location (hsa-mir-199a-3p, hsa-mir-199b-3p, hsa-mir-29b, hsa-mir-101, hsa-mir-320c, hsa-mir-103b), raising the possibility that each genomic location gives rise to only one mature miRNA, which are then perceived as isomiRs.

Primary miRNA transcripts are processed to pre-miRNAs in the nucleus by DROSHA (a dsRNA nuclear type III endoribonuclease) facilitated by DGCR8, and subsequently cleaved in the cytoplasm by DICER to generate functionally mature miRNAs molecules. IsomiRs are generated because both DICER and DROSHA process miRNA precursors imprecisely generating miRNA variants with several plus/minus nucleotides at the 5’ and/or 3’ (41). In addition, miRNAs can also be post-transcriptionally adenylated or uridylated, which could modify miRNA targeting properties and/or their stability (42,43). The importance of endogenously expressed 5’ isomiRs has been demonstrated by Chiang *et al*. (30), where they showed deletion of miR-223 in mouse neutrophils (44), resulted in an increase in expression of predicted targets of a lesser expressed 5’ isomiR of the microRNA. Although it is established 5’ isomiRs can regulate gene expression, the functional reason for their presence is still debated. 5’isomiRs of a given microRNA can have highly over-lapping targets, potentially increasing the specificity of regulation of a particular pathways (24,45). However, 5’ isomiRs can also increase a microRNAs target repertoire (46). Manzano *et al.*, suggest these apparent discrepancies are dependent upon the presence or absence of a U at position 2 of the longer isomiR sequence (47). miR-140-3p.2, the longest 5’ isomiR of miR-140-3p, does not possess a U at position 2, which would suggest miR-140-3p.1 and miR-140-3p.2 have divergent target repertoires, decreasing their effect on any single target (47). Although not directly studied here, 3’isomiRs are widespread, with 3’ adenine additions being most common and reported to decrease the targeting ability of the miRNA (42). Some of the miR-140-3p isomiRs with the same seeds as miR-140-3p.1 and miR-140-3p.2 have 3’ adenine additions, which in combination with divergent 5’isomiRs may account for the poor target repression by miR-140-3p.1 and miR-140-3p.2 compared with miR-140-5p.

Interestingly, there is an inverse correlation between the ability to repress targets (miR-140-5p>miR-140-3p.1>miR-140-3p.2) and expression in cartilage (miR-140-3p.2>miR-140-3p.2>miR-140-5p).

A mutation within miR-140-5p has recently been identified as causing a human skeletal dysplasia (48). Mice with the corresponding miR-140-5p mutation phenocopy the human situation and display a loss of miR-140-5p target repression (in rib chondrocytes). This is thought to contribute to the phenotype since the expression of miR-140-3p.1 and miR-140-3.2 were not affected by the mutation. Interestingly though, the phenotypes of the miR-140-5p mutant mice and *MIR140* null mice show some differences, which could perhaps be accounted for by the latter animals lacking miR-140-3p and the isomiRs described here.

Here we identify discrete targets for miR-140-3p.1 and miR-140-3p.2. LIPA, a cholesterol ester hydrolase, is a target of miR-140-3p.1 but not miR-140-3p.2. *LIPA* expression decreases during chondrocyte development in human (49), consistent with high miR-140-3p.1 expression in chondrocytes. SDC4 (Syndecan-4), a target of miR-140-3p.2 but not miR-140-3p.1, has been implicated in osteoarthritis progression through activation of ADAMTS5 (50), suggesting *SDC4* suppression by miR-140-3p.2 may maintain a healthy cartilage phenotype. *HMGCR* and *HMGCS1* have now been identified as targets of miR-140-3p.1 in breast cancer cells (51), however their expression was not repressed by miR-140-3p.1 in human chondrocytes, indicating miR-140 isomiRs act in a tissue-specific manner. Even where repression was determined, the effects of miR-140-3p.1 and miR-140-3p.2 were relatively small (up to ~25% repression) when compared to the effects of miR-140-5p on a previously published miR-140-5p direct target, FZD6 (~60% repression) (9). Furthermore, contra to the expected miR-140-3p.1 repression of targets, miR-140-3p.1 increased luciferase for the *MARCKSL1*, *KIAA1199* and *MAPILC3A* 3’UTR constructs, possibly suggesting stabilisation of transcript, as previously described for miR-322 and *MEK1* (52).

The target repertoire of miR-140-3p.1 and miR-140-3p.2 are vastly different, due to established miRNA target interaction rules. The only shared binding motif is ‘CTGTGG’, which can be recognised by miR-140-3p.1 as a 6merBS (CTGTGG), or by miR-140-3p.2 as a 7m8-BS or 8mer-BS if followed by T or TA, respectively (CTGTGG**T** or CTGTGG**TA**) (Figure 2A). Indeed the miR-140-3p.1 6merBS (CTGTGG) was the most enriched miR-140-3p.1 motif following miR-140-3p.2 transfection (Figure 3E; red line in bottom right), indicating miR-140-3p isomiRs act according to traditional miRNA targeting rules. Other types of miRNA binding sites such as 3’compensatory binding (53) may be shared between miR-140-3p.1 and miR-140-3p.2, but are less common and more difficult to predict.

Analysis of miRNA target interactions within published skeletal data sets, using the *WWP2* host gene as a surrogate for miRNA expression, along with directly overexpressing the miRNAs and observing changes in gene expression, suggests these types of analyses can improve miRNA target prediction within skeletal tissues. Caution must however be taken as *WWP2* and *MIR140* can display tissue specific expression (35), furthermore, miRNA are often co-expressed with their targets as part of a larger more complex network (54). This type of co-expression may account for the studies where a correlation, rather than an inverse correlation, between *WWP2* and miR-140 targets was observed. Where an inverse correlation was observed between *WWP2* and miR-140 predicted targets this was greatest within cartilage. Together, these data indicate the miR-140-3p isomiR miR-140-3p.1 can function to down-regulate transcript expression *in vitro* and *in vivo*.

In conclusion, analysis of human cartilage sRNA-seq identified an abundant isomiR to miR-140-3p, miR-140-3p.1. Functional analysis in human and murine chondrocytes and analysis of published skeletal datasets suggests the newly identified isomiR is more effective than the original miR-140-3p annotated in miRBase. These data suggest the function of miR-140-3p, which has previously been largely ignored, should be revisited.

## ACCESSION NUMBERS

Microarray data and RNA-seq data are available at NCBI GEO datasets with the accession numbers GSE144374.

## SUPPLEMENTARY DATA

Supplementary Data are available online.

## ACKNOWLEDGEMENT

pLKO.1-puro U6 sgRNA BfuAI large stuffer was a gift from Scot Wolfe (Addgene plasmid # 52628; http://n2t.net/addgene:52628; RRID:Addgene_52628).

## SUPPLEMENTARY FIGURE LEGENDS

**Supplementary figure 1**. Cartilage miRNAs which have abundant 5’ isomiRs. Twenty-nine miRNAs with isomiRs that account for more than 5% of the reads for that miRNA and the number of isomiR reads is greater than 100. miR-1246 isomiR (read count 99) is also shown. 5’ start position of each isomiR relative to the miRBase annotation is shown: the addition of a nucleotide is designated as ‘+1’ and the loss as ‘−1’, ‘0’ represents reads of the miRBase annotation. Histogram indicates read count.

**Supplementary figure 2**. Analysis of published sRNA-seq. miR-140-3p.1 and miR-140-3p.2 were present in multiple sRNA-seq data including; melanoma (Stark *et al*. 2010 (25)), cervix (Witten *et al*. 2010 (26)), lymphocytes (Kuchen *et al*. 2010 (27),) and following immunoprecipitation of Argonaute (AGO) proteins (Burroughs *et al*. (29)). Sequences were designated either miR-140-3p.1 or miR-140-3p.2 based upon seed sequence.

**Supplementary figure 3** Validation of miR-140 null mouse model and rib chondrocyte RNA-seq. (A) Deletion of Mir-140 was attained using the CRISPR/-Cas9 system. The miR-140 locus was targeted in mouse zygotes using two crRNAs (blue line, protospacer adjacent motif (PAM) shown in red). This successfully deleted the 140 locus sequence depicted (KO mir140). (B) Mice were bred to homozygosity with the genotype in wild-type, heterozygous and null mice confirmed through PCR of ear-notch genomic RNA. (C) Total RNA was extracted from rib chondrocytes of postnatal day 7 (P7) mice, reverse transcribed to cDNA and subjected to real time qRT-PCR analysis for gene expression of miR-140-5p and miR-140-3p (canonical). Values were normalised to U6 and plotted as mean ± standard error of the mean (SEM) (number of samples, WT=7, miR-140^−/−^ =6). *** p≤0.001. (D) The RNA from several of the same samples as (C) was subjected to RNA-seq (n=4 for WT and miR-140^−/−^). Principle component analysis of length-scaled TPM (Transcripts Per Million) values segregated WT and KO (blue and red). (E) Heat map of the 1200 most differentially expressed genes.

**Supplementary figure 4** IsomiR read count and enrichment score for all possible sequences derived from miR-140. (A) Read count for all 5’isomiRs encoded from the mouse miR140 stem loop according to miRBase. (B) Sylamer analysis of rib cartilage RNA from mice lacking miR140 stem loop for 7m8 seed binding sites corresponding to every possible miRNA produced from the miR140 locus. Histogram represents average enrichment score for each seed binding motif.

**Supplementary figure 5. Overlap between human and mouse mir140 datasets**. (A) overlap between predicted targets, genes down-regulated following overexpression in human articular chondrocytes (HAC), genes up-regulated in mice lacking the miR140 locus and genes downregulated during human MSC chondrogenesis. (B) Regression analysis of ABCA1 expression and WWP in multiple skeletal tissues, separated based on tissue type.

## SUPPLEMENTARY TABLE LEGENDS

**Supplementary table 1. Cartilage sRNA-seq**. Cartilage sRNA-seq from 3 donors. All sequences including isomiRs aligning to 990 mature miRNAs are shown with read count.

**Supplementary table 2. Predicted targets**. Predicted targets of miR-140-5p, miR-140-3p.1 and miR-140-3p.2 by TargetScanHuman 7.2. Gene lists correspond to Venn diagram in figure 2B.

**Supplementary table 3. Human chondrocyte gene expression**. Human chondrocyte gene expression changes following transfection of miRNA mimics: miR-140-3p.1 vs CON, miR-140-3p.2 vs CON, miR-140-3p.1 vs miR-140-3p.2

**Supplementary table 4. Human chondrocyte gene list**. Gene lists from human chondrocytes: Genes decreased with miR-140-3p.1 vs control, Genes decreased with miR-140-3p.2 vs control, Genes decreased with both miR-140-3p.1 and miR-140-3p.2 vs control, and genes significantly different between miR-140-3p.1 and miR-140-3p.2 when directly compared.

**Supplementary table 5. Human chondrocyte pathway analysis**. Pathway analysis using DAVID for genes decreased with miR-140-3p.1, genes decreased with miR-140-3p.2 genes significantly different between miR-140-3p.1 and miR-140-3p.2.

**Supplementary table 6. MIR140 KO mice gene expression**. RNA-seq of mouse rib cartilage of MIR140 KO mice and WT control mice

**Supplementary table 7. MIR140 KO mice gene lists**. List of up and down-regulated genes in MIR140 KO mice

**Supplementary table 8. MIR140 KO mice pathway analysis**. Pathway analysis using DAVID for genes up-regulated in MIR140 KO mice

**Supplementary table 9. Human vs mouse**. Cross over between predicted targets, human chondrocyte data and mouse KO data

**Supplementary table 10. Oligonucleotides**. Table of oligonucleotides used in this study. Targets of miR-140-3p.1 and miR-140-3p.2 used for cloning into pMiR-GLO by InFusion. Primers used for the generation and genotyping of miR-140^−/−^ mice.

