## Supplementary figures and images for "microRNA-seq of cartilage reveals an over-abundance of miR-140-3p which contains functional isomiRs"

### Supplemental Figures

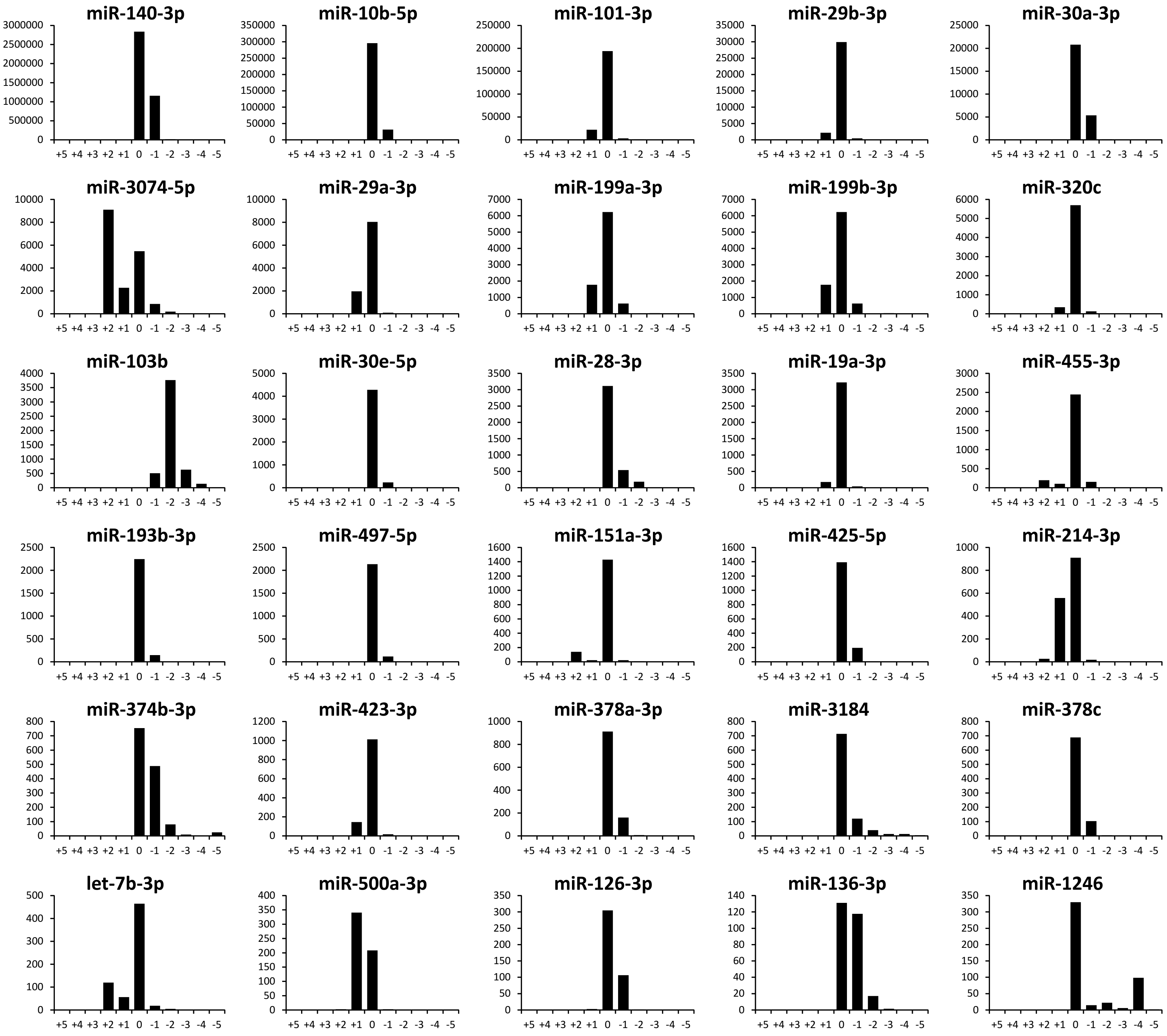

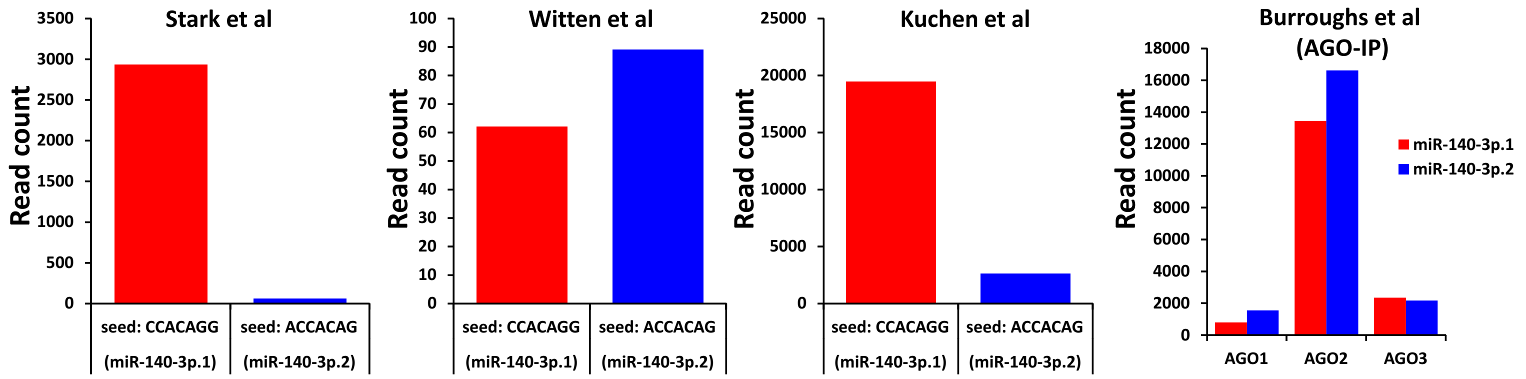

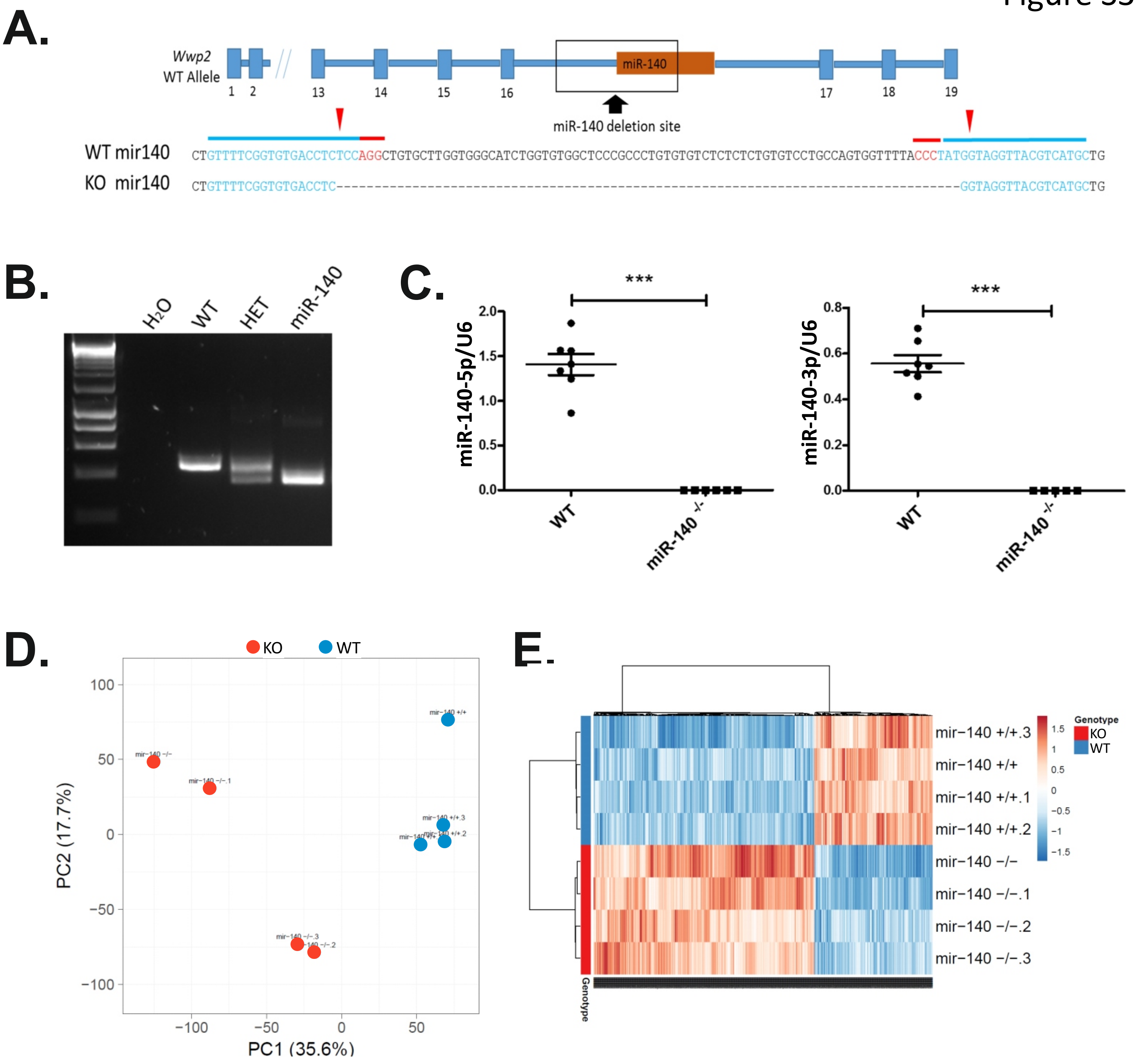

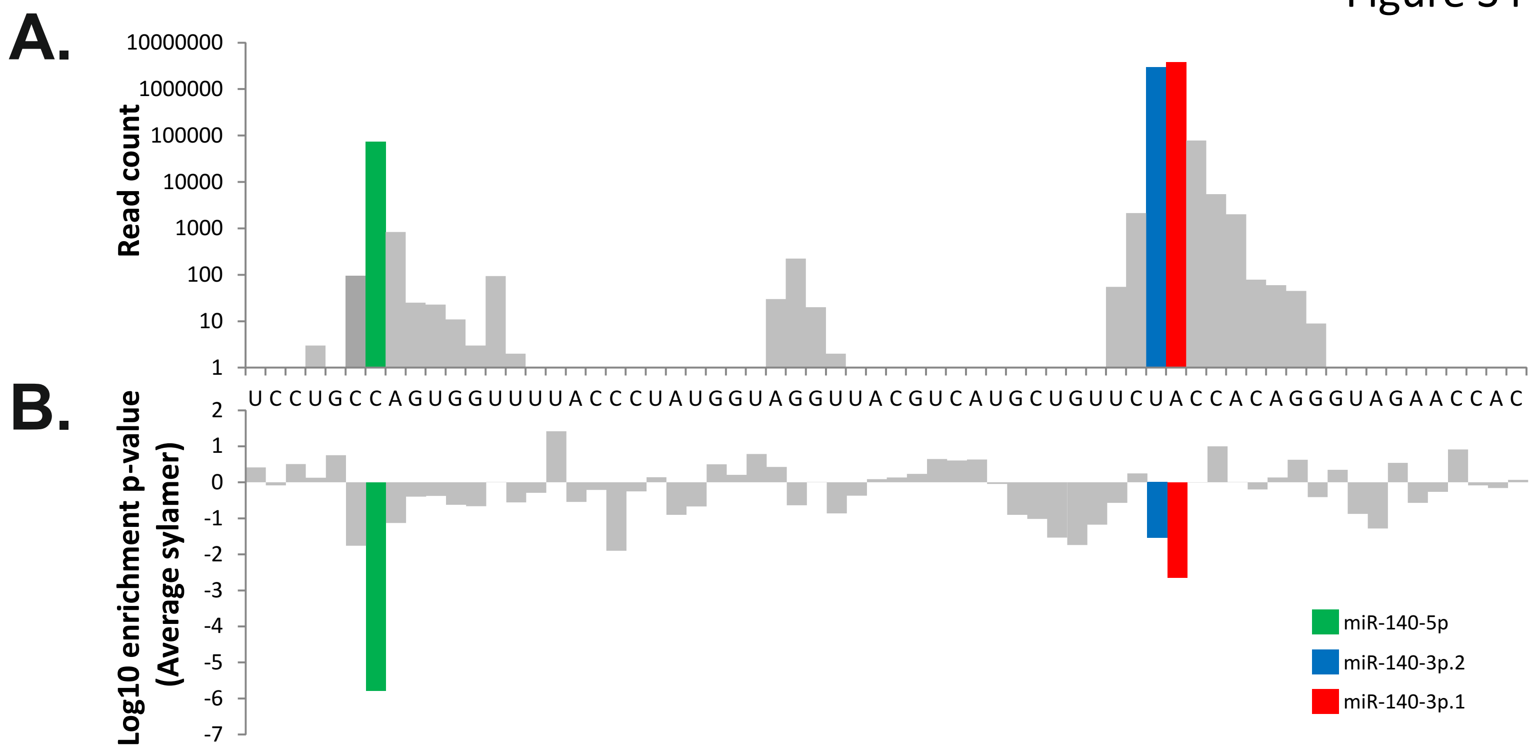

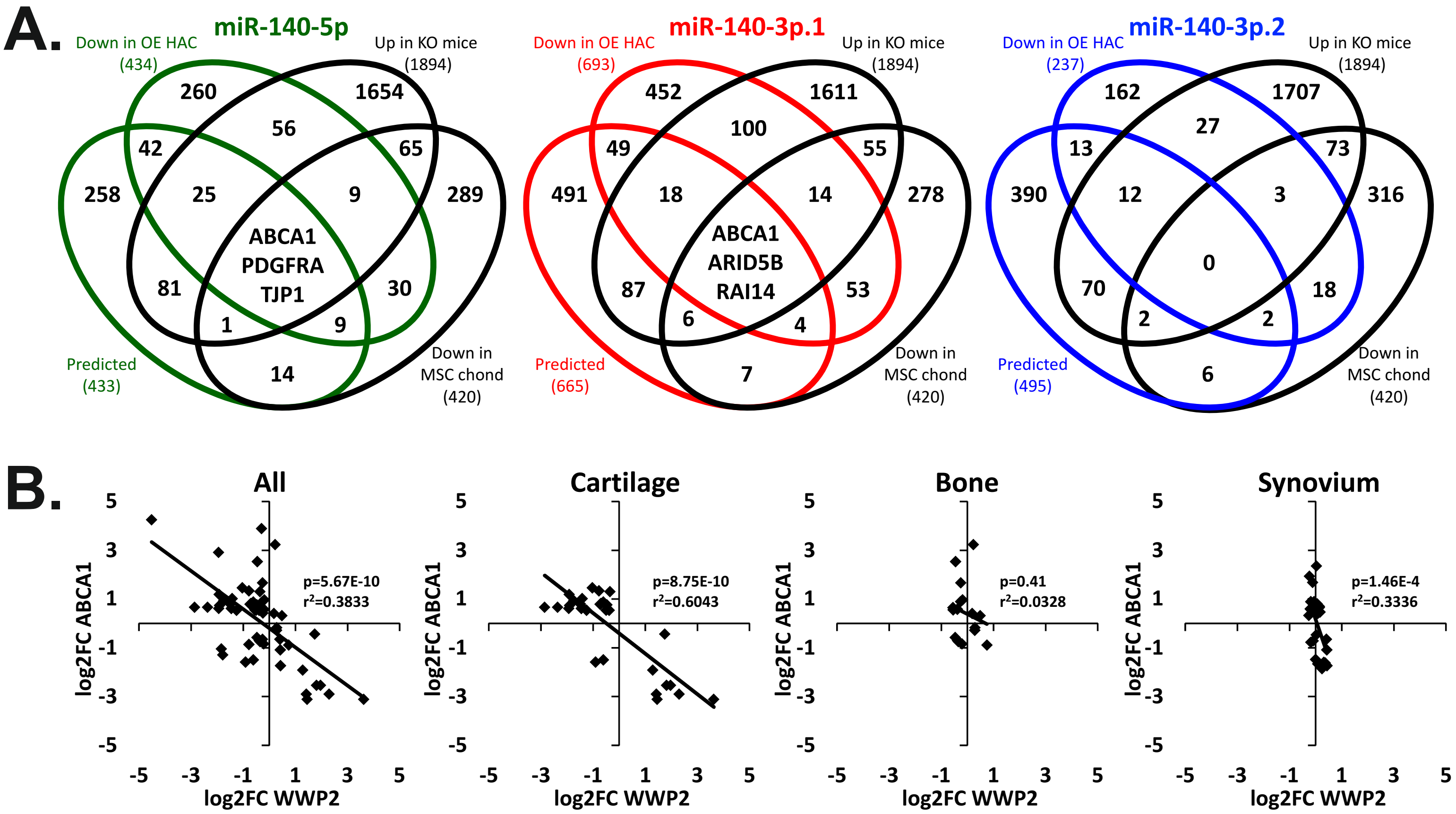
